# Predicting predator-prey interactions in terrestrial endotherms using random forest

**DOI:** 10.1101/2022.09.02.506446

**Authors:** John Llewelyn, Giovanni Strona, Christopher R. Dickman, Aaron C. Greenville, Glenda M. Wardle, Michael S. Y. Lee, Seamus Doherty, Farzin Shabani, Frédérik Saltré, Corey J. A. Bradshaw

## Abstract

Species interactions play a fundamental role in ecosystems. However, few ecological communities have complete data describing such interactions, which is an obstacle to understanding how ecosystems function and respond to perturbations. Because it is often impractical to collect empirical data for all interactions in a community, various methods have been developed to infer interactions. Machine learning is increasingly being used for making interaction predictions, with random forest being one of the most frequently used of these methods. However, performance of random forest in inferring predator-prey interactions in terrestrial vertebrates and its sensitivity to training data quality remain untested. We examined predator-prey interactions in two diverse, primarily terrestrial vertebrate classes: birds and mammals. Combining data from a global interaction dataset and a specific community (Simpson Desert, Australia), we tested how well random forest predicted predator-prey interactions for mammals and birds using species’ ecomorphological and phylogenetic traits. We also tested how variation in training data quality—manipulated by removing records and switching interaction records to non-interactions—affected model performance. We found that random forest could predict predator-prey interactions for birds and mammals using ecomorphological or phylogenetic traits, correctly predicting up to 88% and 67% of interactions and non-interactions in the global and community-specific datasets, respectively. These predictions were accurate even when there were no records in the training data for focal species. In contrast, false non-interactions for focal predators in training data strongly degraded model performance. Our results demonstrate that random forest can identify predator-prey interactions for birds and mammals that have few or no interaction records. Furthermore, our study provides guidance on how to prepare training data to optimise machine-learning classifiers for predicting species interactions, which could help ecologists (*i*) address knowledge gaps and explore network-related questions in data-poor situations, and (*ii*) predict interactions for range-expanding species.

## Introduction

Species interactions are essential for maintaining biodiversity (Thompson 1999) because they determine how energy and nutrients flow among organisms (Traill et al. 2010, Allan et al. 2021) and affect the distribution, survival, and abundance of species (Van der Putten et al. 2010). Species interactions also determine how ecological disturbances—such as those involving invasive species, habitat modification, and climate change—reverberate through ecological communities (Tylianakis et al. 2008, Blois et al. 2013). This is because environmental changes that directly alter one or a subset of species in a community can have flow-on effects for other species via interaction networks (Kaneryd et al. 2012, Morton et al. 2022). Given the need to predict and mitigate human-driven biodiversity declines, interest in modelling interaction networks for biological communities is increasing (Bohan et al. 2017, Strydom et al. 2021; Strona and Bradshaw 2022). However, most species interactions in most ecological communities have not been documented (Hortal et al. 2015, Jordano 2016, Caron et al. 2022). Indeed, it is usually logistically impossible to measure how all species interact in a community—even when only one type of interaction (e.g., predator-prey) is considered (Brousseau et al. 2018, Strona 2022). In addition to the challenge of identifying long-established interactions, the rapid and widespread translocation of species across the planet due to migratory responses to climate change and movement by humans means we also need to identify interactions that could occur between species that previously did not co-occur, and therefore had not previously had the opportunity to interact (Smith and Phillips 2006, Valdovinos et al. 2018).

To overcome gaps in species interaction data, various methods of inferring interactions using species traits/proxies can be applied, such as quantile-regression trophic niche-space modelling (Gravel et al. 2013, Llewelyn et al. 2021), Bayesian linear modelling (Caron et al. 2022), and several machine-learning approaches (e.g., *K* nearest-neighbour—(Desjardins-Proulx et al. 2017, Pichler et al. 2020), neural networks/deep learning methods—(Fricke et al. 2022), random forests—(Sydenham et al. 2021), and ensemble methods—(Poisot 2022)). Of these, random forest is a popular method due to its ability to learn nonlinear interactions between predictor variables and predict interactions accurately (Desjardins-Proulx et al. 2017, Laigle et al. 2018, Sydenham et al. 2021). Random forests have been used to predict various interaction types, including plant-pollinator (Ornai and Keasar 2020, Sydenham et al. 2021), parasite-host (Kotula et al. 2020), and predator-prey interactions (Desjardins-Proulx et al. 2017, Parravicini et al. 2020). They have also been used to identify which species’ traits are most important for predicting whether species interact (Dellinger et al. 2019, Klomberg et al. 2022). Using training data consisting of interacting and non-interacting species, random forests can learn how ecomorphological traits, phylogenetic position, and/or taxonomy (as a proxy for phylogenetic position; Laigle et al. 2018) relate to which species interact. However, questions remain regarding the ability of random forests to predict different types of interactions in different taxonomic groups, which species’ data/traits are best for inferring interactions, and how to fine-tune training datasets (Poisot 2022). Furthermore, investigating ways to optimise training data for predicting species interactions using random forest will likely help guide procedures for optimising other machine learning methods.

Identifying predator-prey interactions is central to community ecology because these interactions influence both top-down and bottom-up processes such as energy transfer and extinction cascades (Sinclair et al. 2003). Random forests have been applied to predict such interactions, but these applications have largely been limited to fish communities (Parravicini et al. 2020, Strona et al. 2021) and soil invertebrates (Desjardins-Proulx et al. 2017, Laigle et al. 2018). Indeed, the performance of random forests for inferring predator-prey interactions among terrestrial vertebrates has not been assessed despite this group including many species of high conservation concern (Tilman et al. 2017, Cox et al. 2022) and of important ecological function (e.g., ecological engineers; Samson and Knopf 1996).

If we are to model terrestrial ecosystems realistically to predict and minimise the risk of extinction cascades, reliable inference of trophic interactions is essential. However, trophic-related traits and network structure differ among terrestrial vertebrates, fish, and invertebrates (Liem 1984, Liem 1990, Shurin and Gruner 2006, Miller-ter Kuile et al. 2022), suggesting traits and methods (such as random forest and other machine learning techniques) that effectively predict interactions in fish and invertebrates might not necessarily work well for terrestrial vertebrates. These vertebrates are diverse structurally, phylogenetically, and functionally, potentially making the ‘rules’ determining interactions complex and difficult for machine-learning classifiers to identify (whereas just one trait—body size—largely determines trophic interactions in fish due to their gape limitation; Gravel et al. 2013, Morales-Castilla et al. 2015). Furthermore, interaction data for many terrestrial vertebrate species have been recorded opportunistically or not at all (although detailed diet data do exist for some species; Pringle and Hutchinson 2020), potentially leading to sparse (i.e., missing most prey for most predators) and taxonomically biased datasets for training and testing random forests. Having sparse interaction records for predators in training data is problematic because machine learning classifiers such as random forests need to be trained with data that include both interacting (e.g., predator and prey) and non-interacting species pairs, with non-interacting species pairs usually assigned by assuming that undocumented interactions imply no interaction (i.e., pseudo-non-interactions generated from presence-only data; Pichler et al. 2020). When applied to incomplete (sparse) datasets, this process can identify undocumented but truly interacting species pairs as non-interacting pairs (but see Strona et al. 2021). Importantly, such ‘false non-interactions’ in training data likely reduce the reliability of interactions and non-interactions inferred from random forests and other classifiers. Similarly, random forests and other classifiers trained on datasets that are taxonomically biased might perform poorly when making predictions for species not represented in the training data (Strydom et al. 2022).

We designed analyses to test whether random forest could accurately predict predator-prey interactions within and between two diverse terrestrial vertebrate classes—birds and mammals—using both a global (*GloBI*) and a single-community (Simpson Desert, Australia) predator-prey interaction dataset. We also assessed (*i*) the importance of ecomorphological traits and phylogenetically based latent traits (i.e., as proxies for phylogenetically conserved traits associated with trophic interactions; Benandi et al. 2022) for inference accuracy, (*ii*) whether prediction accuracy improved by removing predators from the training dataset that had few interaction records (i.e., those with scarce diet records and were, therefore, prone to generating false non-interactions), and (*iii*) the sensitivity of model performance to the quality of the training data. To test sensitivity to the quality of training data, we compared two scenarios that could occur as a result of sparse training data: (a) a reduction in the number of interaction and non-interaction records used to train the models, and (b) a reduction in the number of interaction records and an increase in the number of false non-interaction records used to train the models. We also investigated how these changes affected model performance when they were restricted to different components of the training dataset (i.e., testing the effect of taxonomic bias on performance).

Our results confirm that: (*i*) random forests can predict interactions in birds and mammals, (*ii*) models trained on interaction datasets including either ecomorphological traits or phylogenetic position (or both) can perform well, (*iii*) removing predators from the training data that have few recorded interactions therein can improve prediction accuracy, (*iv*) models can identify the prey of predators that have no interaction records in the training data, and (*v*) model performance is especially sensitive to false non-interactions in the training data that involve records of predatory species for which predictions are being made (i.e., focal predators). Our results highlight the unrealised potential for the application of random forests for predicting interactions among terrestrial vertebrates. Furthermore, they show to which components of the training data random forests are most sensitive, thereby identifying the most efficient ways to increase accuracy via fine-tuning of training data—highlighting steps that would likely also improve species-interaction inference by machine learning classifiers more generally.

## Methods

We used two interaction datasets combined with ecomorphological and phylogenetic information to test the performance of random forests for predicting predator-prey interactions in birds and mammals (Supporting information Figure S1 shows flowchart of dataset preparation). We also tested how the performance of these models was influenced by the type of species data used, the filtering of training data by removing predators with few interaction records in that data, and the quality of the training data. The two interaction datasets were: (1) a dataset from the ***Glo****bal **B**iotic **I**nteractions database* (globalbioticinteractions.org; Poelen et al. 2014) supplemented with interaction data from diet studies done in Australia (Llewelyn 2022), and (2) an ecosystem-specific dataset from the Simpson Desert in Australia that focuses on seven predators for which detailed dietary information is available (Llewelyn 2022). We focused on birds and mammals rather than on all tetrapods due to the availability of detailed trait (Wilman et al. 2014) and phylogenetic (vertlife.org) information for birds and mammals (detailed trait databases for reptiles and amphibians are far from complete), thereby providing a wide range of traits for predicting interactions.

## Datasets

### Global data

We extracted all predator-prey interactions involving birds and mammals from *GloBI* (accessed 29/05/2020 and 26/06/2020 for birds and mammals as consumers, respectively). However, *GloBI* contains few records of species found in Oceania (including Australia) relative to other regions of the world: of the 3329 unique species combinations in the records we extracted from *GloBI*, only 109 involved a predator and prey species both found in Oceania. Supervised learners, including random forests, need to be trained on data that capture the variation present in the species for which predictions are being made, and so geographic and taxonomic biases in training data could reduce prediction accuracy. We therefore added interactions involving Simpson Desert species and their relatives to the *GloBI* data (Supporting information). We added data for these species by merging two additional datasets with *GloBI*: (*i*) predator-prey records from the Simpson Desert that did not involve the seven focal predators (focal predator = species for which we made predictions), and (*ii*) interactions of the focal Simpson Desert predators with non-Simpson Desert prey elsewhere in Australia (i.e., from diet studies of the focal predators outside the Simpson Desert; no interactions between the focal predators and Simpson Desert species were included in the global/training data). In both cases, we built these datasets by aggregating predator-prey records from diet studies (Supporting information). We then combined these additional interaction datasets with the *GloBI* data (adding 363 records) to create an enhanced global interaction dataset (Llewelyn 2022).

To predict species interactions using random forests, we required a training dataset that included interacting and non-interacting pairs of species. Like most interaction datasets, our global dataset did not include non-interactions i.e., since relationships are based on presence-only data. We therefore generated pseudo-non-interactions (‘non-interactions’ henceforth) by taking all possible two-species combinations from the species in our global interaction dataset and removing those combinations documented as ‘interacting’ (supporting information). We also removed combinations where either the ‘predator’ does not consume vertebrates (e.g., strictly herbivorous species could not be potential predators) or the combination involved species not found on the same continent (using the `rgbif` package in R; Chamberlain et al. 2022). We removed these non-interactions because they would not help models predict which sympatric species known predators preyed upon (i.e., the type of interactions we aimed to predict in the Simpson Desert). Although the non-interactions we generated would have included many sympatric, truly non-interacting species combinations, they would have also included false non-interactions (species combination assigned as not interacting, but that do indeed interact) as well as allopatric species pairs that would interact if they were sympatric.

Random forests use explanatory variables such as species’ traits to learn and predict associations (Desjardins-Proulx et al. 2017). We therefore added trait data to all the predator and prey species in the interaction and non-interaction datasets (Supporting information). These traits included: body mass, time of activity (whether the species displays nocturnal, diurnal, and/or crepuscular activity), ability to fly, resource groups consumed (plants, vertebrates, and/or invertebrates), and percentage use of different habitat strata (8 habitat categories for birds and 5 for mammals; Wilman et al. 2014; Supporting information). We added phylogenetic information by calculating eigenvectors describing a species’ position in the phylogeny (eigenvector mapping; Guénard et al. 2013), a potential proxy for explanatory variables which are phylogenetically conserved. We extracted phylogenies from VertLife.org, and used the `MPSEM` (Guénard and Legendre 2022) and `ape` (Paradis and Schliep 2019 p. 0) R packages to handle phylogenetic data and extract eigenvectors.

We also identified which species pairs in the non-interaction data included a potential prey species from within the predator’s preferred prey-size range (Supporting information). We did this so the ratio of non-interaction records from inside *versus* outside the preferred size range in training datasets could be adjusted to optimise performance of random forests. This is analogous to optimising species distribution models by adjusting the ratio of pseudo-absences sampled from among *versus* outside observed occurrences (Barbet-Massin et al. 2012). To identify a preferred size range, we log-transformed (log*_e_*) predator and prey body masses to linearise body-size relationships (Gravel et al. 2013). We then resampled the interaction dataset to create 100 training and testing dataset pairs using all 3692 interaction records in each of the 100 pairs (75:25 interaction records in training:testing datasets).

To each training set we fitted a series of upper and lower quantile regressions capturing the middle 0.99 to 0.8 quantiles at 0.01 quantile increments. We included a prey mass × predator taxonomic class interaction in the regression models to allow different body-size relationships for predatory birds and mammals (Llewelyn et al. 2021). We then combined each of the 100 testing sets with an equal number of species pairs randomly sampled from the non-interaction set (also with body mass log*_e_*-transformed), and applied the quantile regression models (i.e., upper and lower quantile regressions combinations) built in the previous step to the testing sets. Species combinations in the testing sets that included a potential prey species from within the upper and lower quantile ranges derived from the regressions were predicted as prey, and those outside these limits were excluded as prey.

We assessed model performance by comparing predictions to documented interactions (i.e., asking whether the quantile regression models correctly identified which combinations came from the interaction and non-interaction datasets), and used these results to calculate the true skill statistic to identify the optimal quantile regression model for assigning predator-prey interactions. The true skill statistic is a metric that can effectively evaluate the performance of classifiers such as those predicting species interactions (Poisot 2022). A true skill statistic = 1 indicates a model that perfectly predicts interactions and non-interactions, whereas when it is 0, the model performs no better than assigning links randomly. The model capturing the middle 0.84 quantiles performed best (true skill statistic = 0.26), so we used that model to classify species pairs in the non-interaction dataset as those that included a potential prey from within the predator’s preferred size range and those that included a potential prey from outside the predator’s preferred size range.

### Simpson Desert testing data

It is unclear how the accuracy of random forests trained and tested on data including species from many different communities (such as was the case for our global datasets) translates to model performance when applied to single ecological communities (Parravicini et al. 2020). We therefore tested model performance when applied to seven common predators from the Simpson Desert, a community we chose because it has been intensively studied for over 30 years and has detailed information on species composition and the diets of some predators. We used the species documented at established field sites in the Simpson Desert as the species assemblage (i.e., potential prey), rather than all species documented in the Simpson Desert bioregion, because the former includes sympatric species that have the potential to interact whereas the latter would likely include species that seldom have the opportunity to interact.

The Simpson Desert bird and mammal assemblage from these sites includes 64 vertebrate-consuming bird and mammal species (49 birds and 15 mammals). However, the diets of many of these species have not been studied in detail. When testing the performance of random forests for predicting predator-prey interactions, we therefore restricted the Simpson Desert testing dataset to seven predatory species with detailed dietary information from studies both in the Simpson Desert and other arid regions in Australia. These seven focal predators included four birds—*Aquila audax*, *Falco berigora*, *Hamirostra melanosternon*, *Tyto alba*, and three mammals—*Canis dingo*, *Felis catus*, *Vulpes vulpes*. *Felis catus* (domestic cats) and *Vulpes vulpes* (red fox) are recent introductions to Australia (since European colonisation in 1788), *Canis dingo* (dingo) was brought to Australia by people > 3000 years ago (Fillios et al. 2012), and the remaining focal predators naturally occur in Australia.

To construct the dataset, we aggregated data from published and unpublished diet studies (excluding species consumed as carrion) of the focal predators in the Simpson Desert (Supporting information; Llewelyn 2022). We also included records from diet studies done outside the Simpson Desert involving species pairs found in the Simpson Desert (i.e., we assumed that if trophic interactions occurred between the species in other communities, they would also occur in the Simpson Desert; Llewelyn 2022), and records involving the focal predators and Simpson Desert species in the global interaction dataset (we removed these records from the global dataset). None of the studies we used to build the Simpson Desert interaction dataset reported non-interactions. We therefore generated non-interactions by assuming any species pairs involving the focal predator and the species recorded from the Simpson Desert field sites (157 bird and mammal species) that have not been documented as predator and prey are not predator and prey. We added the same trait and phylogenetic information to the Simpson Desert data as described for the global dataset. However, we did not include a column indicating if non-interacting species pairs in the Simpson Desert data involved potential prey from within the predator’s preferred size range because we only used the Simpson Desert dataset to test model performance (i.e., we did not need to optimise the sampling of these non-interactions because they were not used for training random forests).

## Modelling

We used the `ranger` package (Wright and Ziegler 2017) in R and the Flinders University high performance computer (Flinders 2021) to apply random forests using the model-based recursive partitioning algorithm (Breiman 2001). In the first phase, we optimised the random forest models and tested how performance was affected by (a) the number and choice of variables included in the random forests, and (b) removing predators that had few documented interactions in the training data (i.e., those whose diet is likely not adequately captured in the data). In the second phase, we tested how changes to the quality of the training dataset—including removing records and increasing the number of false non-interactions—influenced model performance (Supporting information shows flowchart of modelling steps and what different models were used for). When optimising models, we assessed model performance by comparing predictions to documented interactions and used these results to calculate the true skill statistic (we also calculated other performance metrics to confirm our results; see Supporting information). A true skill statistic of 1 indicates a model that perfectly predicts interactions and non-interactions, a score > 0.75 is ‘excellent’, between 0.4 to 0.75 is ‘fair to good’, and < 0.4 indicates ‘poor’ performance (Fleiss et al.2003).

### 1. Model optimisation and variables

We used the global dataset and cross-validation to optimise training datasets and model parameters, and we compared performance of models trained using: (*i*) only ecomorphological traits, (*ii*) only phylogenetic eigenvectors, or (*iii*) both types of data (Supporting information). We also compared performance of models that incorporated many *versus* few explanatory variables/traits to test whether more variables resulted in higher accuracy or overfitting. There were 42 variables in the many-variable ecomorphological model (21 each for predator and prey; Llewelyn 2022) because there were 21 ecomorphological variables in the databases we built (Supporting information). To be consistent, we used 42 phylogenetic eigenvectors in the many-variable phylogenetic model (first 21 phylogenetic eigenvectors for predator and prey). In the few-variable models, we included 10 variables (a number we chose arbitrarily that is substantially smaller than the 42 variables in the many-variable models). These 10 variables were the most important ecomorphological or phylogenetic variables (5 each for predator and prey) for predicting interactions, as determined by permutation, which measures the impact of changing a variable’s value on model accuracy. To identify the most important variables, we sequentially removed the least-important variable and repeated the permutation and removal steps until 10 variables remained (retained variables for each model listed in Supporting information; Llewelyn 2022). Although a variable’s importance in predicting interactions does not imply causation (Arif and MacNeil 2022), knowing which traits are most important for predicting interactions can help researchers avoid spending time and resources collecting data on traits that are uninformative. The many-and few-variable models that incorporated both types of data simply combined the variables from the respective ecomorphological and phylogenetic models (i.e., either 84 or 20 variables in total).

We also tested whether performance was enhanced or eroded by removing records from the global dataset involving predators with few (< 5) interaction records. We posited that predators that had only a small proportion of their diet documented in the dataset could bias results and lead to the generation of many false non-interactions because, for example, models might be unduly influenced by predators that had not been studied in detail – rather than matching of predator and prey traits. Removing records involving these predators reduced the number of interactions from 3,692 in the original global dataset to 3,424 in the modified global dataset, and reduced the number of non-interactions from 454,924 to 89,184.

To optimise training datasets and model parameters, we divided the global dataset’s interactions (original and modified data) into training and testing datasets (ratio: 75:25 training:testing), and combined them with non-interaction data. The number of non-interaction records added to the testing datasets equalled the number of interaction records in that dataset, whereas we varied the number of non-interactions in the training datasets to identify the optimal (highest true skill statistic) ratio. We replicated the process of generating training and testing datasets each time we applied a random forest in the parameter-optimization and cross-validation steps described below.

We weighted non-interactions such that their combined weight was equal to that of the observed interactions because models built with weighted non-interactions performed slightly better than those built using unweighted non-interactions (Supporting information; similar to results from species distribution modelling; Barbet-Massin et al. 2012). These weightings determine the probability an interaction/non-interaction is selected in the bootstrap samples for the trees built from the training datasets. We optimised the training datasets and random forests in terms of (*i*) number of non-interactions in the training data (see above), (*ii*) the ratio of non-interactions from inside *versus* outside preferred size range in the training data, (*iii*) the probability threshold for assigning interactions—because random forests can estimate probabilities for binary responses such as interact *versus* not interact (Malley et al. 2012), (*iv*) the number of variables randomly sampled as candidates at each split in the trees, (*v*) the number of decision trees in the random forest, and (*vi*) the maximum depth of the trees.

We optimised these parameters by iterating through values for each parameter individually, running 5 random forests at each value (on 5 resampled training and testing datasets) and calculating the mean true skill statistic for models at each value. We identified the parameter values that gave models with the highest true skill statistic. We separately optimised the 12 models that differed in terms of included variables (i.e., the many- *versus* few-variable ecomorphological, phylogenetic, and combined data models) and whether the original or modified global dataset was used (Supporting information shows optimised parameter sets). We then compared performance of these optimised models by applying them to 100 training and testing datasets generated using the global dataset cross-validation design described above, and calculated the true skill statistic for each of these 100 cases.

After comparing performance of the twelve models with the global dataset, we tested their performance when trained on the global dataset (with hyper-parameters set to optimise performance, as identified in the previous step) and applied to the Simpson Desert data (Supporting information). Again, we trained and applied 100 random forests for each of the twelve models, and assessed their performance with the true skill statistic. We resampled non-interactions for the training dataset in these 100 analyses because there were more non-interactions than needed, whereas we used all available interaction records in the global datasets (3,692 and 3,424 for the original and modified global datasets, respectively) in each of the 100 training datasets.

### 2. Quality of training dataset and model performance

To test the influence of the size of the training dataset, taxonomic coverage, and prevalence of false non-interactions (true interactions classified as non-interactions) on model performance, we reran the analyses with the optimised model that performed best on the Simpson Desert data (i.e., the many ecomorphological and phylogenetic variable model trained on the modified global dataset; Figure 1f; Supporting information) but varied the number of records, taxonomic coverage, and prevalence of false non-interactions in the training datasets (Supporting information).

**Figure 1.**
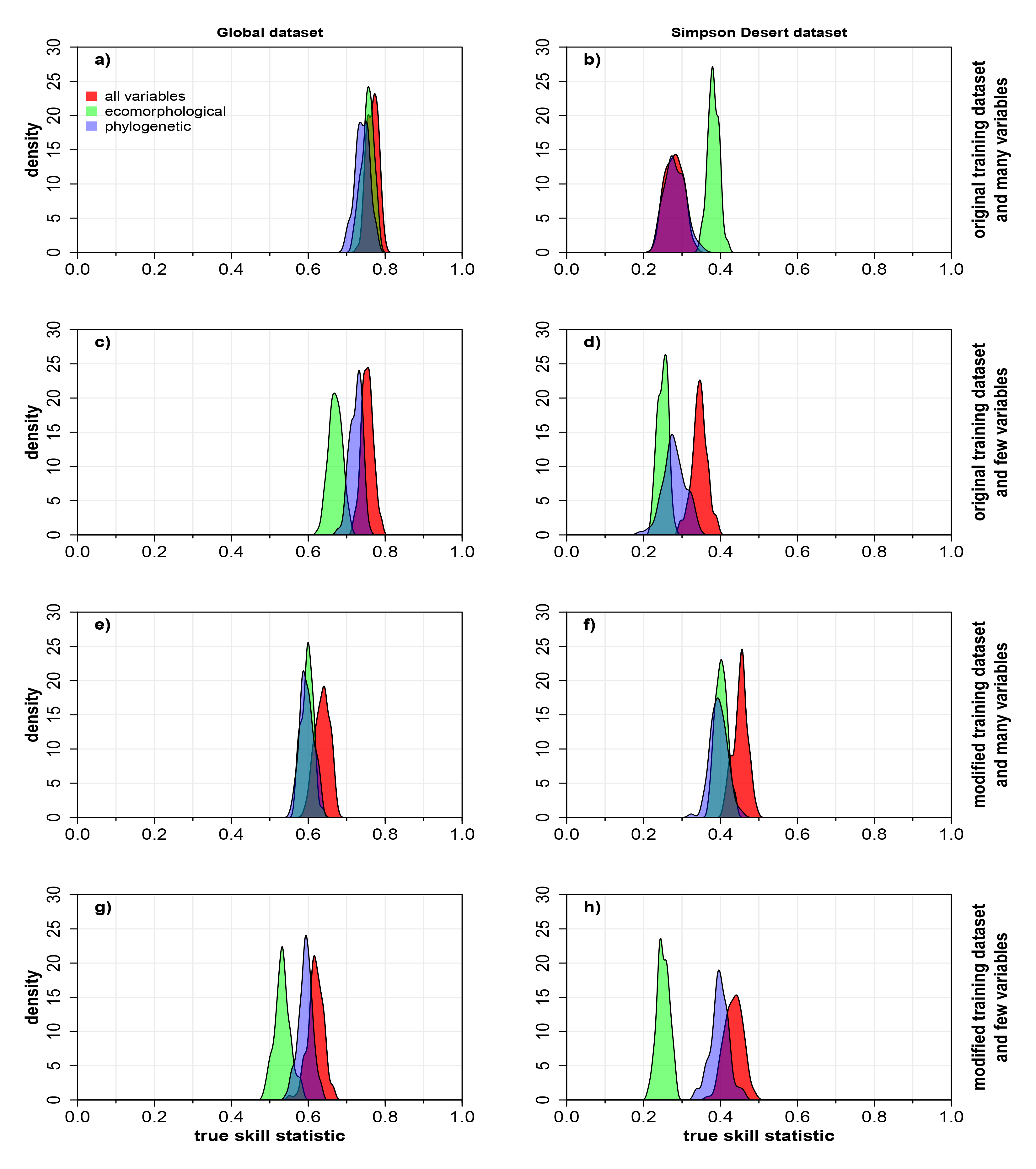
Density plots comparing the performance (true skill statistic) of random forest models predicting predator-prey interactions in birds and mammals. The random forests used either ecomorphological traits, phylogenetic eigenvectors, or both types of data, and were applied to two interaction datasets: a global interaction dataset (panels in the left column) and a specific ecosystem dataset (7 predators and their potential prey/sympatric species in the Simpson Desert; panels in the right column). We also assessed the effects of using many variables (42, 42, and 84 variables, for ecomorphological, phylogenetic eigenvector, and both data-type models) or few variables (10, 10, and 20 variables, for ecomorphological, phylogenetic eigenvector, and both data-type models), and restricting the training dataset to predators with ≥ 5 prey records (modified training dataset). Panels (a) and (b) show results from models that use the complete training dataset and many variables; (c) and (d) show results from models that use the complete training dataset and few variables; (e) and (f) show results from models that use the modified training dataset and many variables; and (g) and (h) show results from models that use the modified training dataset and few variables.

We deliberately reduced the quality of training data in eight ways to test specific sensitivity hypotheses. We modified the entire training dataset by (*i*) randomly removing records and (*ii*) randomly switching interaction records to non-interactions (false non-interactions). We hypothesised that model performance would be sensitive to removal of training data because having more records helps the models learn associations between species traits and trophic links, and we hypothesised that models would be sensitive to false non-interactions because random forests would learn the wrong association when trained with erroneous data. We also tested what impact data removal and false non-interactions had when they were restricted to subsets of the training data that included either: (*iii*, *iv*) focal prey, (*v*, *vi*) focal predators, or (*vii*, *viii*) interactions where neither prey nor predators were focal/Simpson Desert species. We hypothesised that model performance would be especially sensitive to changes involving focal predators and prey because these records involve the species for which predictions are made. Here, the associations between these species’ traits and trophic interactions should be especially important for the model to learn. Furthermore, we hypothesised that model performance would be sensitive, albeit less so, to removing records and increasing the proportion of false non-interactions in non-Simpson Desert (non-focal) interactions because these interactions teach the random forests trait-trophic associations that also apply to Simpson Desert species. Assessing the sensitivity to changes in the quality of training data involving the focal species is important because it assesses the suitability of random forests for inferring interactions of species that have few/no interaction records, a common phenomenon.

For each of the eight ways training dataset quality was reduced, we made these changes at 1% increments relative to the initial number of records in the modified group (e.g., 1% increments of the total number of interactions involving the focal predator), generating 10 training datasets at each increment. We trained the random forests on these reduced-quality datasets, applied them to the Simpson Desert data, and calculated true skill statistics 10 times at each increment. We plotted results to make visual comparison of the sensitivity of model performance to each of the eight types of artificial variation in the quality of training data, and we compared model performance when all relevant data were modified to quantify the impacts of each type of modification.

We also examined the mechanisms underlying changes in model performance by testing how removing records or increasing false non-interactions for the focal predator influenced the: (*i*) relative probability (or suitability) of interacting assigned by the random forest to each potential prey for each predator (i.e., asking whether changes to the training dataset affected which species were identified as the most likely prey), and (*ii*) mean probability assigned to potential prey for each predator (i.e., asking whether changes to the training dataset cause an overall increase or decrease in the probabilities assigned to potential prey). If lower model performance reflects a systematic shift in all the probabilities (upward or downward), rather than a change in which potential prey are identified as the most likely, the models could still be used to predict a predator’s prey because the rank order of probability of different prey remains similar irrespective of absolute model performance and absolute probability.

To test for changes in relative prey suitability, we measured the correlation (Pearson’s *r*) between prey probabilities assigned by models trained on reduced-quality training datasets with that assigned by models trained on the full training dataset (Supporting information). To test for changes in mean probability assigned to potential prey overall, we subtracted the mean probability assigned to potential prey for each predator as calculated by the full training-data models from that calculated by the models trained on the reduced-quality training datasets (Supporting information). For both analyses, we reduced the quality of the training dataset at 1% increments (10 replicate training datasets/models at each increment) and plotted the results to compare visually. We hypothesised that removing records and including more false non-interactions in the training dataset would disrupt the assignment of relative suitability and reduce assigned probabilities overall.

## Results

### Variables and model performance

Random forests trained and tested on the global data had mean true skill statistics ranging from 0.53 to 0.77 (Figure 1a,c,e,g), indicating ‘good’ to ‘excellent’ performance (Fleiss et al. 2003). These models tended to perform better when we used the original global interaction dataset rather than the modified dataset that had predators with few prey records removed (mean difference in true skill statistic between models = 0.14; Figure 1a and Figure 1c *versus* 1e and 1g). Using few or many variables in the phylogenetic or phylogenetic/ecomorphological variable models had little effect on performance when applied to the global datasets (difference in mean true skill statistic < 0.02 in all cases; Figure 1), but there was a slight decrease in performance associated with using few variables in the ecomorphological-only models (difference in mean true skill statistic = 0.08 and 0.06 between the many-and few-variable ecomorphological models when the original or modified global dataset was used, respectively; Figure 1a *versus* 1c, Figure 1e *versus* 1g).

The models trained on the global dataset and applied to the Simpson Desert data yielded mean true skill statistics between 0.25 and 0.45, indicating ‘poor’ to ‘fair’ performance (Fleiss et al. 2003; Figure 1). Those that used both phylogenetic and ecomorphological variables performed well when trained on the modified global dataset, yielding mean true skill statistics of 0.43 ± 0.02 (mean ± standard deviation) and 0.45 ± 0.02 when we used few or many variables, respectively (Figure 1f,h). Using few *versus* many variables did not strongly affect the performance of the phylogenetic-only or phylogenetic-and ecomorphological-variable models (difference in mean true skill statistic < 0.07 in all four cases; Figure 1). In contrast, the few-variable ecomorphological models did not perform as well as their many-variable counterparts (difference in mean true skill statistic > 0.13 between the many-and few-variable ecomorphological models when using either the original or modified global datasets Figure 1b *versus* 1d, Figure 1f *versus* 1h).

Using the model that performed best on the Simpson Desert data (i.e., the many-variable model with phylogenetic and ecomorphological traits trained on the modified global dataset; Figure 1f), performance predicting interaction for recently introduced predators—cats and foxes—was better than that for native predators (true skill statistic = 0.56 ± 0.03 and 0.41 ± 0.02 for recently introduced and native predators, respectively; Figure 2).

**Figure 2.**
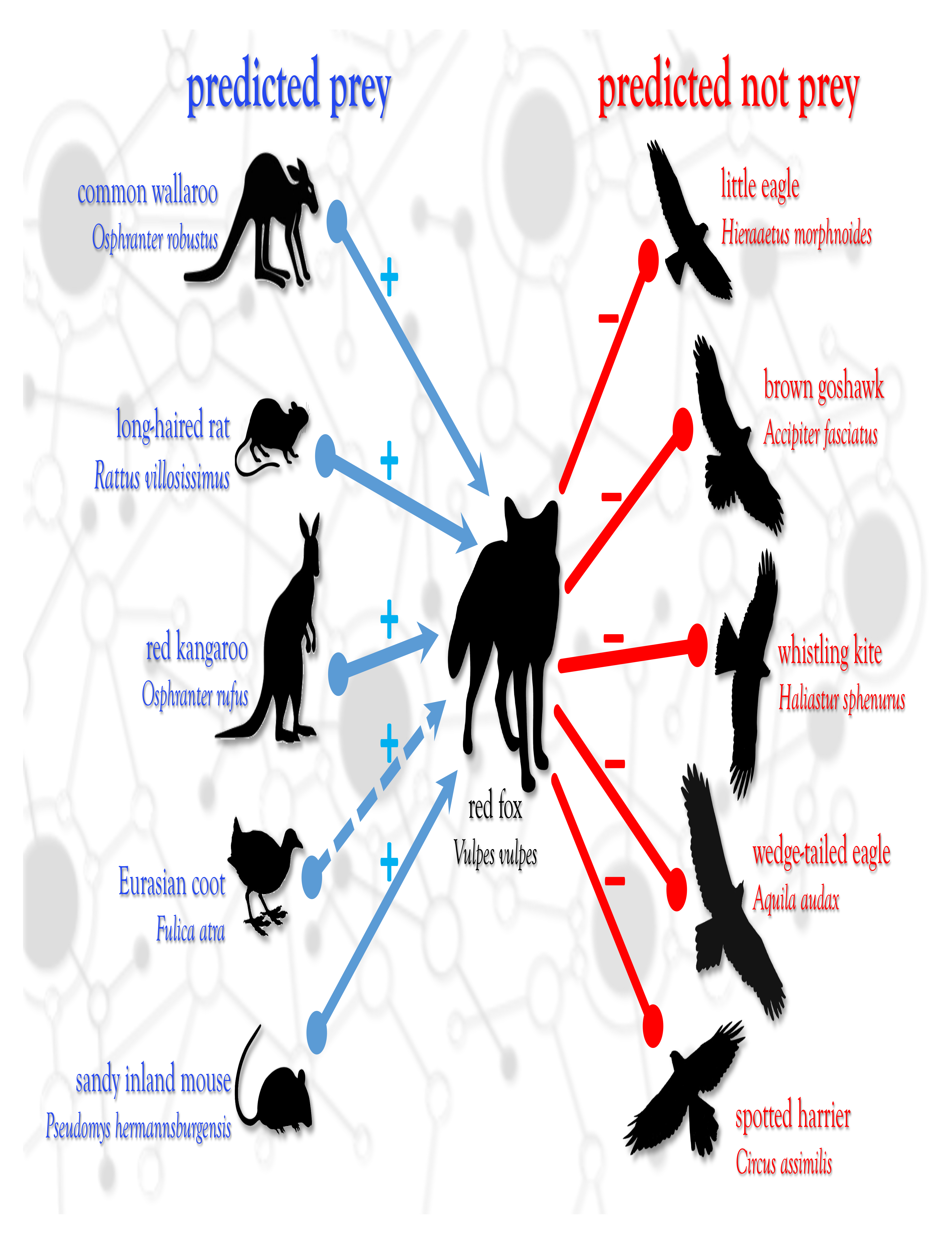
The five most-and least-likely mammalian and bird prey for the red fox (*Vulpes vulpes*) in the Simpson Desert predicted by random forest. Solid blue arrows indicate prey that were predicted and observed, dashed blue arrows show prey that were predicted but not observed (potentially indicating interactions that occur but have not yet been recorded), and red lines indicate species that were not predicted or observed as prey. The five most-likely prey according to the many-variable phylogenetic and ecomorphological model (run 100 times and trained on the modified training data) were the common wallaroo (*Osphranter robustus*), long-haired rat (*Rattus villosissimus*), red kangaroo (*Osphranter rufus*), Eurasian coot (*Fulica atra*), and sandy inland mouse (*Pseudomys hermannsburgensis*). The five least-likely prey were the little eagle (*Hieraaetus morphnoides*), brown goshawk (*Accipiter fasciatus*), whistling kite (*Haliastur sphenurus*), wedge-tailed eagle (*Aquila audax*), and spotted harrier (*Circus assimilis*).

### Effects of training dataset size, false non-interactions, and taxonomic coverage

We altered the global (modified) training dataset by randomly removing records and changing interaction records to false non-interactions, applying these changes both to the entire training dataset as well as to three subsets: (*i*) records involving the focal prey (i.e., species found in the Simpson Desert), (*ii*) records involving the focal predators (i.e., the 7 Simpson Desert predators for which we predicted interactions), and (*iii*) records involving species not found in the Simpson Desert. Removing records reduced performance when modifying either the entire training dataset (Figure 3a; true skill statistic reduced to 0 when all records removed because random forests cannot run without training data) or the focal-predator component only (Figure 3e; true skill statistic reduced to 0.24 ± 0.01 [mean ± standard deviation] when all focal-predator records were removed). However, these effects became pronounced only when > 80% of the interaction and non-interaction records were removed (Figure 3a,e), which equates to removing > 15,750 or 1,240 records from the entire or focal-predator components of the training datasets, respectively. Removing records involving the focal prey or non-Simpson Desert species had only weak or no effect on model performance (Figure 3c,g; true skill static = 0.36 ± 0.02 and 0.46 ± 0.01 when removing all focal prey or all non-Simpson Desert species, respectively).

**Figure 3.**
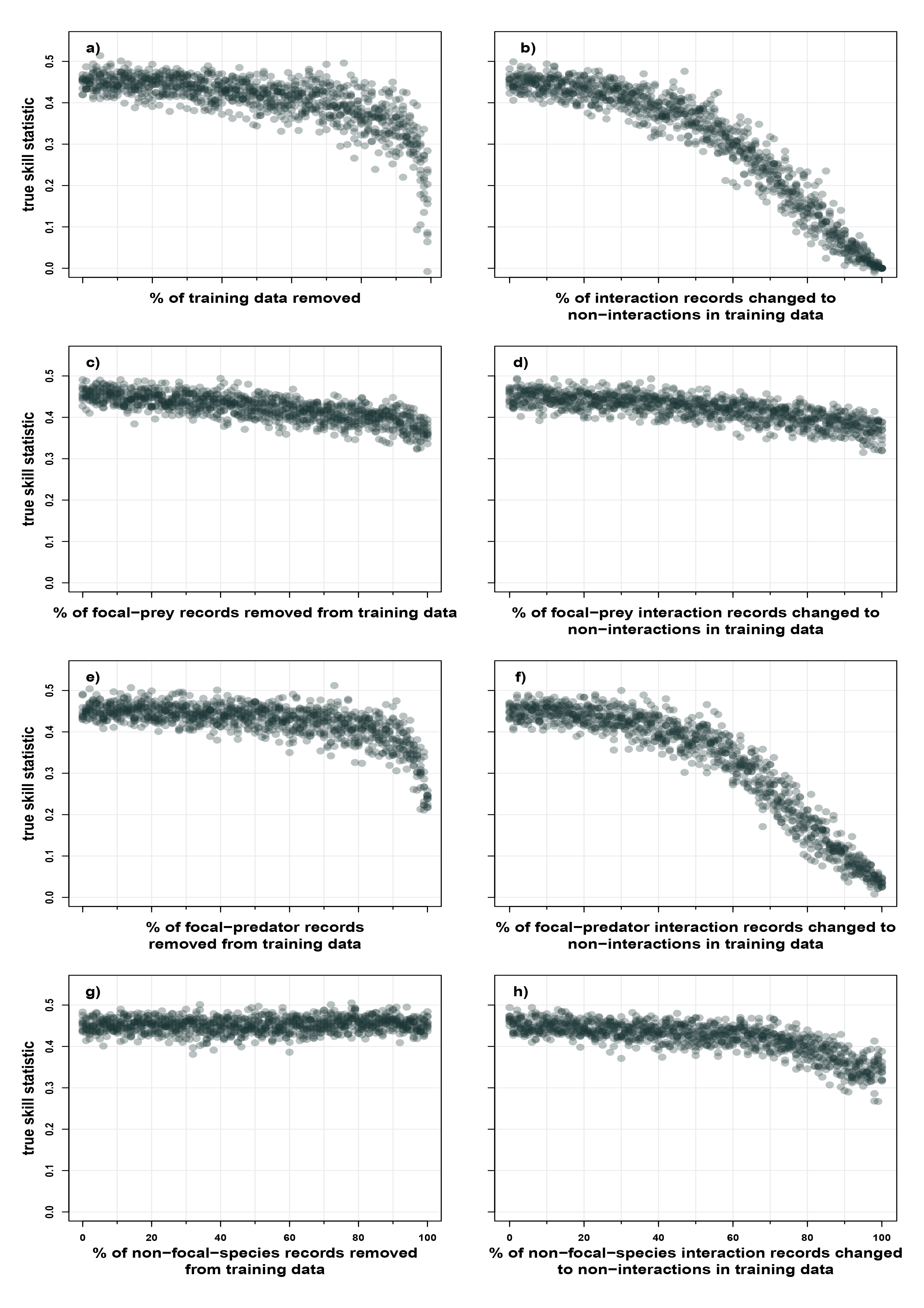
Effect of training-dataset quality on the performance of random forest for predicting predator-prey interactions in birds and mammals. Each panel shows performance when modifying a percentage of either the whole or specific taxonomic groups in the training dataset. Panels (a) and (b) show performance when changing the whole dataset, panels (c) and (d) show performance when restricting changes to focal prey, panels (e) and (f) show performance when restricting changes to focal predators, and panels (g) and (h) show performance when restricting changes to non-focal (non-Simpson Desert) species. Panels in the left column show the effects of removing records from the training data, whereas panels in the right column show the effects of switching interaction records to non-interactions (false non-interactions). We trained random forests on the modified global predator-prey interaction dataset (i.e., restricted to predators with ≥ 5 prey records) and applied them to seven predators and their potential prey from the Simpson Desert. Plot points indicate performance (according to the true skill statistic) when applied to the Simpson Desert predators.

The effect of false non-interactions on model performance was stronger than that of removing records. False non-interactions assigned at random to interaction records in either the entire training dataset or the component of the training dataset involving focal predators degraded model performance more rapidly (Figure 3b,f; true skill static reduced to 0 and 0.04 ± 0.01 for models when changing all records or all records involving focal predators to non-interactions, respectively). These declines were triggered when changing > 40% of the interaction records in the relevant group to false non-interactions (Figure 3b,f), which equates to 1,370 records in the entire training-dataset analysis and 235 in the focal-predator analysis. False non-interactions in the training data involving the focal prey (Figure 3d) or non-Simpson Desert species (Figure 3h) caused slight declines in model performance (true skill static reduced to 0.36 ± 0.03 and 0.34 ± 0.02 when changing all interactions records to false non-interactions for focal prey species and non-Simpson Desert species, respectively).

### Focal-predator training data and the mechanisms underlying changes in model performance

The correlation between probabilities assigned to potential prey by models trained on the full *versus* record-removed global datasets showed a gradual decrease as more interactions involving the focal predator were removed from the training data, and this decline became more rapid when removing > 80% of records (Figure 4a). The correlation between these probabilities reduced to 0.59 ± 0.02 when all records involving focal predators were removed from the training dataset, indicating that models identified many of the same species as prey even when there were no records involving the focal predator in the training data. The mean probability assigned to potential prey for each predator was reduced by removing records of the focal predator from the training data and, like the correlation results, this change was small until 80% of records were removed (Figure 4c). After removing all records, the probabilities assigned to potential prey were on average 0.17 ± 0.01 lower compared to those assigned by models trained on the full dataset.

**Figure 4.**
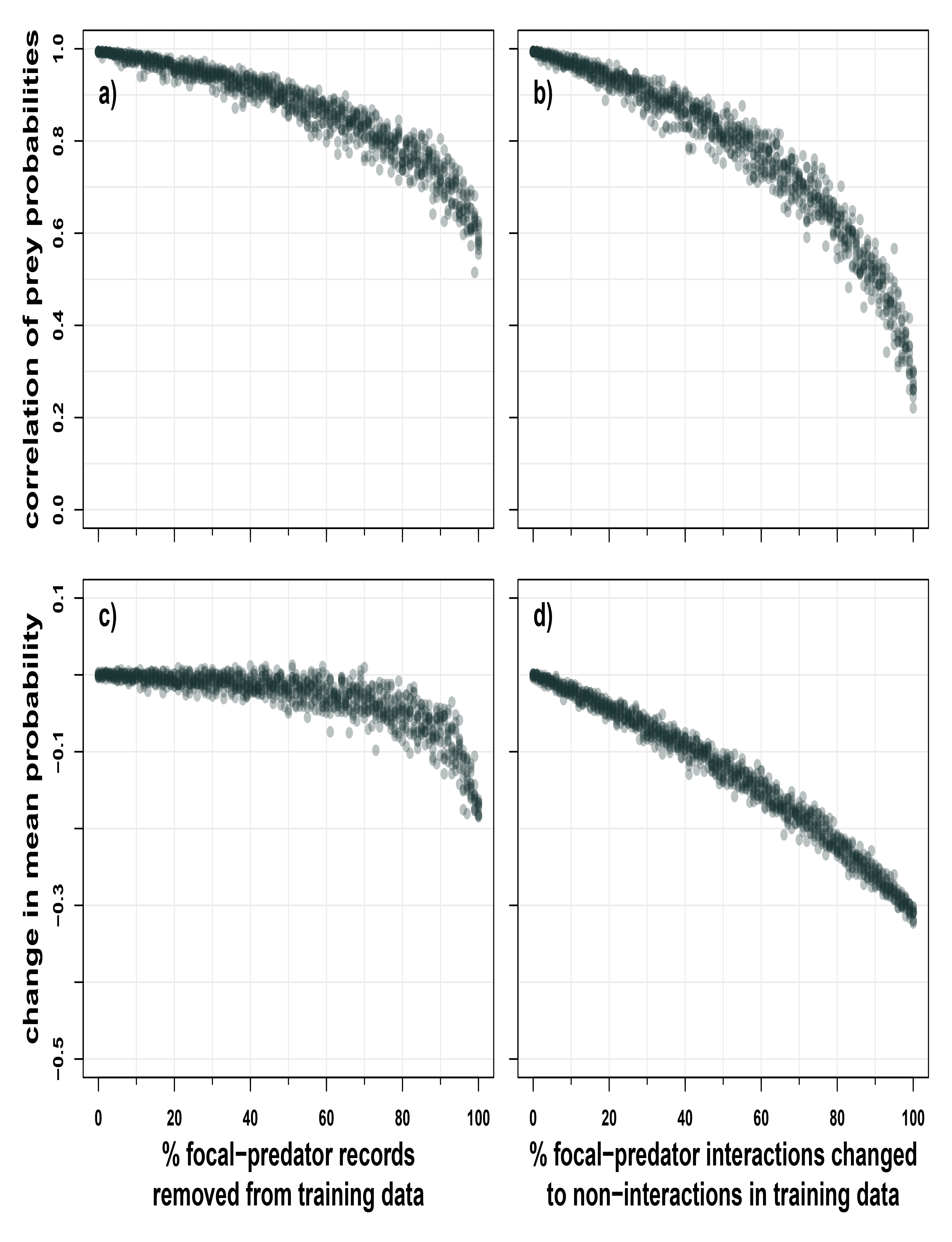
Effects of the quality of the training dataset on (*i*) the predicted relative suitability of prey and (*ii*) the mean probabilities assigned to potential prey by random forests. Panels in the left column show the effects of reducing the number of interactions records involving the focal predator in the training dataset, whereas panels in the right column show the effect of switching interaction records involving the focal predator to non-interactions (false non-interactions). Panels (a) and (b) show the correlation between prey probabilities assigned by the models trained on the changed datasets with that assigned by models trained on the complete (modified) training dataset. Panels (c) and (d) show the mean probability (suitability) assigned to potential prey by models trained on the changed datasets minus those values assigned by the models trained on the complete (modified) training datasets. We trained these random forests on the modified global predator-prey interaction dataset (restricted to predators with ≥ 5 prey records) and applied them to seven predators and their potential prey from the Simpson Desert.

Both the correlation and change in mean probability assigned to potential prey were more sensitive to false non-interactions involving the focal predators than to removing their records from the training dataset, showing more rapid declines (Figure 4a *versus* 4b, Figure 4c *versus* 4d). When changing all interactions involving the focal predators in the training data to non-interactions, the correlation between probabilities assigned to prey decreased to 0.27 ± 0.03 (Figure 4b) and the mean probabilities assigned to potential prey dropped by 0.31 ± 0.01 (Figure 4d).

## Discussion

Our results demonstrate that random forests can predict the prey of predatory birds and mammals even when few or no interaction records involving the focal predator are included in training data (Figure 4a). Applying random forests to predict predator-prey interactions for these predators (and terrestrial vertebrates more generally) to facilitate network modelling of terrestrial vertebrate systems is therefore supported (Brousseau et al. 2018). However, our analyses highlight that caution is needed in applying the random forest and other machine-learning approaches depending on the quality, quantity, and type of training data available, and in the face of these limitations we suggest that filtering training data could improve model performance (Figure 5).

**Figure 5.**
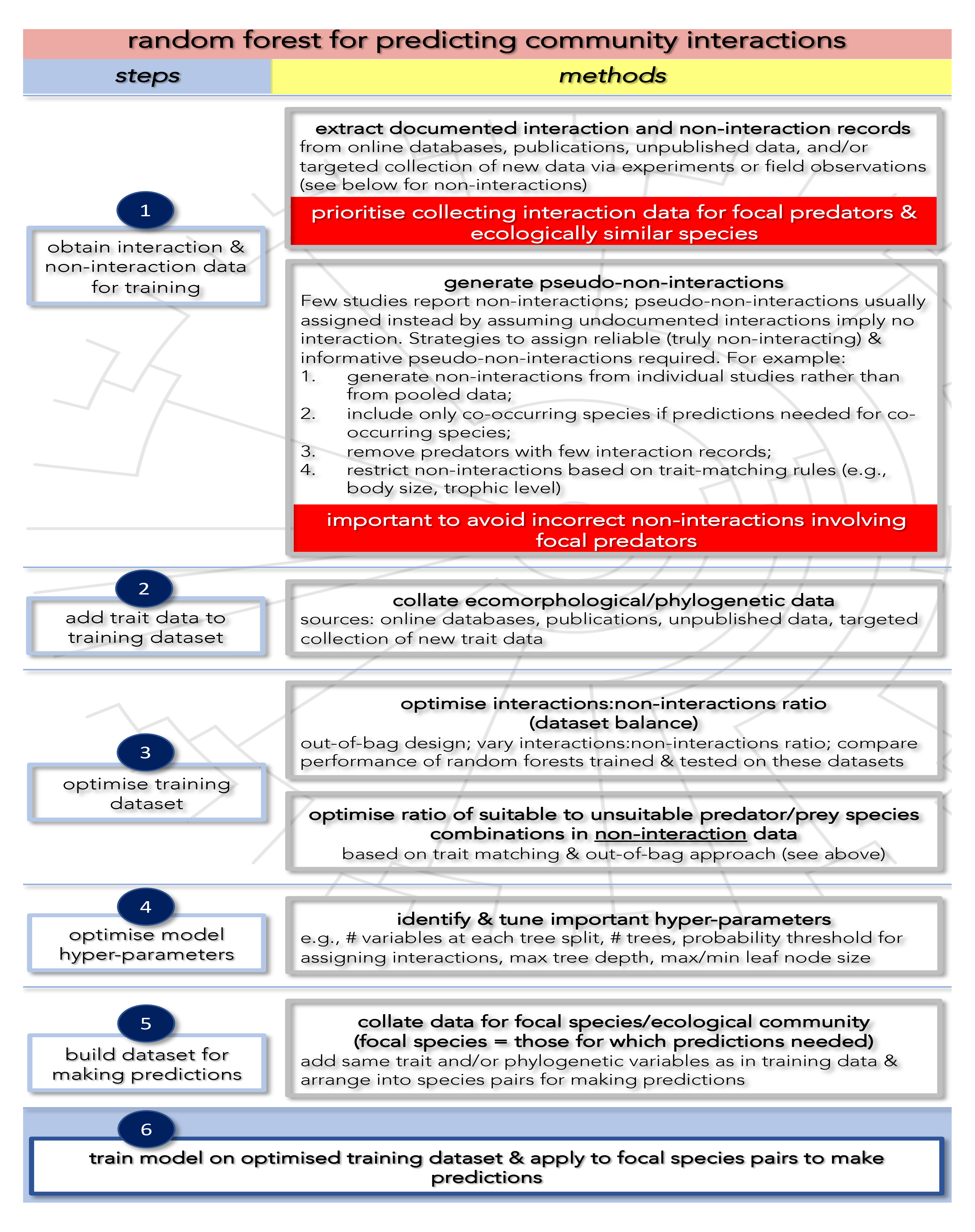
Framework for preparing and optimising random forest models for predicting species interactions. Most of these steps and methods are also relevant to the application of other machine-learning approaches for inferring species interactions.

We found that random forests could predict bird and mammal interactions in both our global and ecosystem-specific datasets (including for native and recently introduced predators). Models based on ecomorphological, phylogenetic, or both types of traits, and those using few or many traits, could perform similarly well (Figure 1)—consistent with previous studies showing that either data type can be used for predicting interactions in other taxa (Faisal et al. 2010, Morales-Castilla et al. 2015, Brousseau et al. 2018, Pomeranz et al. 2019, Elmasri et al. 2020) and that few traits/variables are required to make accurate predictions (Desjardins-Proulx et al. 2017). Because phylogenetic information is available for most terrestrial vertebrates (e.g., VertLife.org), ecological/morphological trait databases are increasingly common and comprehensive (Jones et al. 2009, Oliveira et al. 2017, e.g., Herberstein et al. 2022), and interaction data for many species are available from open-source databases (e.g., globalbioticinteractions.org; web-of-life.es) and publications, random forests and other machine learning methods can now easily be applied to such data to predict predator-prey interactions for most vertebrate communities, and aid in the ecological network and invasive species modelling required to guide conservation decisions.

We found that model performance was robust to the random removal of records from the training dataset. In fact, we had to remove > 80% of records (>15,750 individual records) before an obvious change in performance occurred (Figure 3a). Thus, models that use large training datasets are unlikely to be sensitive to random additions or removal of interaction and non-interaction records.

The removal of records involving focal predators only caused a prominent decline in model performance when we removed > 80% (Figure 3e) of records in that group, but this translates to fewer (1,240) records in total needing to be removed (as compared to the random removal of records). Conversely, the complete removal of records involving focal-prey species or non-focal species had little impact on model performance (Figure 3c,g). Thus, declines in model performance in the random-removal analyses were largely driven by the loss of focal predator records. This reduction in model performance following removal of focal-predator records was due to every prey species being assigned consistently lower probabilities and a slight shuffling of prey preference (Figure 4a,c). Because *relative* prey preferences were largely maintained even when all training records involving the focal predator were removed from training data (mean Pearson *r* = 0.59; Figure 4a), our results suggest prey species could be predicted for predators that do not have any interaction data, which is likely the case for many understudied, rare, or extinct species. The ability to predict trophic interactions for extinct species accurately has important applications for understanding past extinction events (Ripple and Van Valkenburgh 2010, Pires et al. 2015, Llewelyn et al. 2021), while the capacity to predict interactions for rare species could help identify cryptic factors that threaten their persistence (Smith and Phillips 2006, Doherty et al. 2016). However, additional work to assess the requirements of training data for predicting interactions for species without interaction records is still required. For example, it is unclear how phylogenetically and/or ecologically similar species in the training data need to be to the focal species for models to be able to predict trophic interactions accurately.

In contrast to removal, false non-interactions in the training data reduced model performance more rapidly, triggering lower performance when > 40% of the interaction records were changed to non-interactions. This sensitivity highlights an important issue: most studies generate non-interactions by assuming that undocumented interactions imply true non-interaction; however, this assumption could generate many false non-interactions, especially if interaction records are sparse (i.e., if many of the predators in the training data have only a small proportion of their interactions documented). To avoid generating false non-interactions, one can restrict non-interactions in training datasets to species combinations where the predator is too small to eat the potential prey and/or the potential predator comes from a low trophic level (Strona et al. 2021). However, applying this approach could reduce the capacity of random forests to identify unsuitable prey from within a predator’s preferred prey-size range or for high trophic-level species. Indeed, the optimal ratio of non-interactions sampled from within *versus* outside preferred size range in the model that performed best on our Simpson Desert data (i.e., the many-variable, both data-types model) was 5:2 (Supporting information), suggesting that non-interactions of suitably-sized potential prey should be included in training data.

Alternatively, filtering training data by removing predators that have few interaction records could help avoid false non-interactions because many of the interactions/prey of these predators are likely missing (i.e., they have sparse interaction records, although these species would also include some predators with narrow diet breadths). When applied to the Simpson Desert data, our models trained on the global dataset modified by removing predators with < 5 interaction records performed better than those trained on the original/full global dataset (Figure 3). This result might reflect a reduction of false non-interactions in the modified training dataset. Thus, removing predators/consumers that are missing most of their interactions from training datasets could be an effective way to filter training data and improve model performance.

Model performance was especially sensitive to false non-interactions in the training data involving focal predators. This type of change in data quality disrupted relative prey preferences and lowered probabilities assigned to interactions more extensively than did reducing the number of records involving focal predators (Figure 4a *versus* 4b, Figure 4c *versus* 4d), reinforcing the importance of avoiding false non-interactions—especially for focal predators—when generating pseudo-non-interactions for training data; otherwise, such data can change which species are identified as most-likely prey. Although removing predators/consumers with few interaction records from training data and/or restricting the generation of non-interactions based on relative size and trophic level are two simple methods for avoiding false non-interactions as we discussed, there are other strategies that could be applied to the entire-or focal-predator-component of training datasets to minimise the number of false non-interactions. For example, false non-interactions could be minimised by generating non-interactions on a case-by-case basis (similar to the ‘target-group absences’ approach used in species distribution modelling; Barbet-Massin et al. 2012), an approach shown to improve the accuracy of predator-prey interactions predicted by deep-learning models (Fricke et al. 2022). We generated non-interactions for the global dataset using interaction data from different regions, studies, and sources, but this approach could generate false non-interactions, such as when pairs of species are allopatric in studies that contributed to the training data and so are designated as non-interacting, but the component species interact when/where they are sympatric. This approach could also generate misleading non-interactions, such as pairs of currently allopatric species that might interact should future conditions force them to become sympatric. Instead, training data non-interactions could be generated based on individual studies or locations of recorded interactions describing what a predator does and does not eat in a particular area. Such a nuanced approach would identify more reliable and informative non-interactions than would be achieved by assuming that any undocumented pairs from an incomplete global database are truly not predator and prey.

Our models performed better with the global dataset than with the Simpson Desert data. This difference might indicate that the random forests learnt patterns in the global dataset that are not as effective when re-applied to specific communities. For example, the random forests might have learnt from the global dataset that predators with particular traits do not eat prey with particular traits, but predators or prey with those traits might not be present in the Simpson Desert. In other words, learning the relationship did not help the random forest predict interactions in the Simpson Desert. Alternatively, random forests might learn which predators had few or many prey records within the global dataset—another pattern that would be less effective at identifying interactions and non-interactions when applied to the Simpson Desert predators due to the similarity among focal predators in terms of number of interactions in the training data (60% of predators in the original/full global dataset had < 5 interaction records whereas all Simpson Desert focal predators had > 10 interaction records in the global dataset). This might explain why our models did not perform as well on the global data when we used the modified dataset rather than the original/full global dataset for training and testing, because we had removed predators with few interaction records from the modified dataset. Geographic biases in the global training dataset could also lead to weaker performance on our ecosystem-specific dataset (Tuia et al. 2022). Irrespective of what caused the difference in model performance, our results indicate that performance based on global datasets might not be a strong indicator of performance when applied to single communities.

Previous studies applying random forests in aquatic and invertebrate communities have tended to report higher performance than our models achieved (mean true skill statistic consistently > 0.6; Desjardins-Proulx et al. 2017, Laigle et al. 2018, Brousseau et al. 2018, Parravicini et al. 2020, Strona et al. 2021). There are several possible explanations for this discrepancy, including previous studies (*i*) pooling prey species into broad resource categories (Parravicini et al. 2020), (*ii*) using higher-quality training data (Laigle et al. 2018, Parravicini et al. 2020), (*iii*) including resource predictions for a wider range of trophic groups, including easily-excluded non-interactions such as herbivores and/or detritivores eating other animals (Desjardins-Proulx et al. 2017), and (*iv*) focusing on taxa whose interactions might be easier to predict than for terrestrial vertebrates (although further testing is required to evaluate this possibility). Another potential means of improving the performance of random forests that neither we nor others have yet addressed is to include detail on intraspecific variation in traits and interactions, including ontogenetic shifts (González-Varo and Traveset 2016).

Random forest and other machine learning approaches are useful and flexible tools that have great potential in terms of predicting species interactions. While global interaction databases are a valuable resource for training and testing such models, they currently include only a small fraction of real-world species interactions. Furthermore, for the consumers that are included in such databases, in most cases only a small subset of their trophic interactions are documented, thereby increasing the likelihood of generating false-non-interactions. Databases therefore need to be expanded, and a framework for generating non-interactions that avoids or minimises false non-interactions and retains informative non-interactions is required (Figure 5 shows a proposed framework). By making such changes, the application of random forests and other machine-learning methods (including ensemble approaches) can be further honed to infer species interactions more accurately.

