## Supporting information for "Predicting predator-prey interactions in terrestrial endotherms using random forest"

**Contents**

| **Item** | **Page** | **Object** |
|---|---|---|
| Figure S1 | 1 | data-preparation flowchart |
| Figure S2 | 2 | modelling flowchart |
| Table S1 | 3 | ecomorphological variables |
| Table S2 | 4 | datasets/models and their application |
| Table S3 | 5 | performance (several metrics) of different models when applied to global and Simpson Desert datasets |
| Table S4 | 6 | variables retained in the few-variable models |
| Table S5 | 7 | preliminary analysis comparing performance of models with interactions:non-interactions weighted *versus* unweighted in the training data |
| Table S6 | 8 | optimised parameters for different models |

Fig. S1

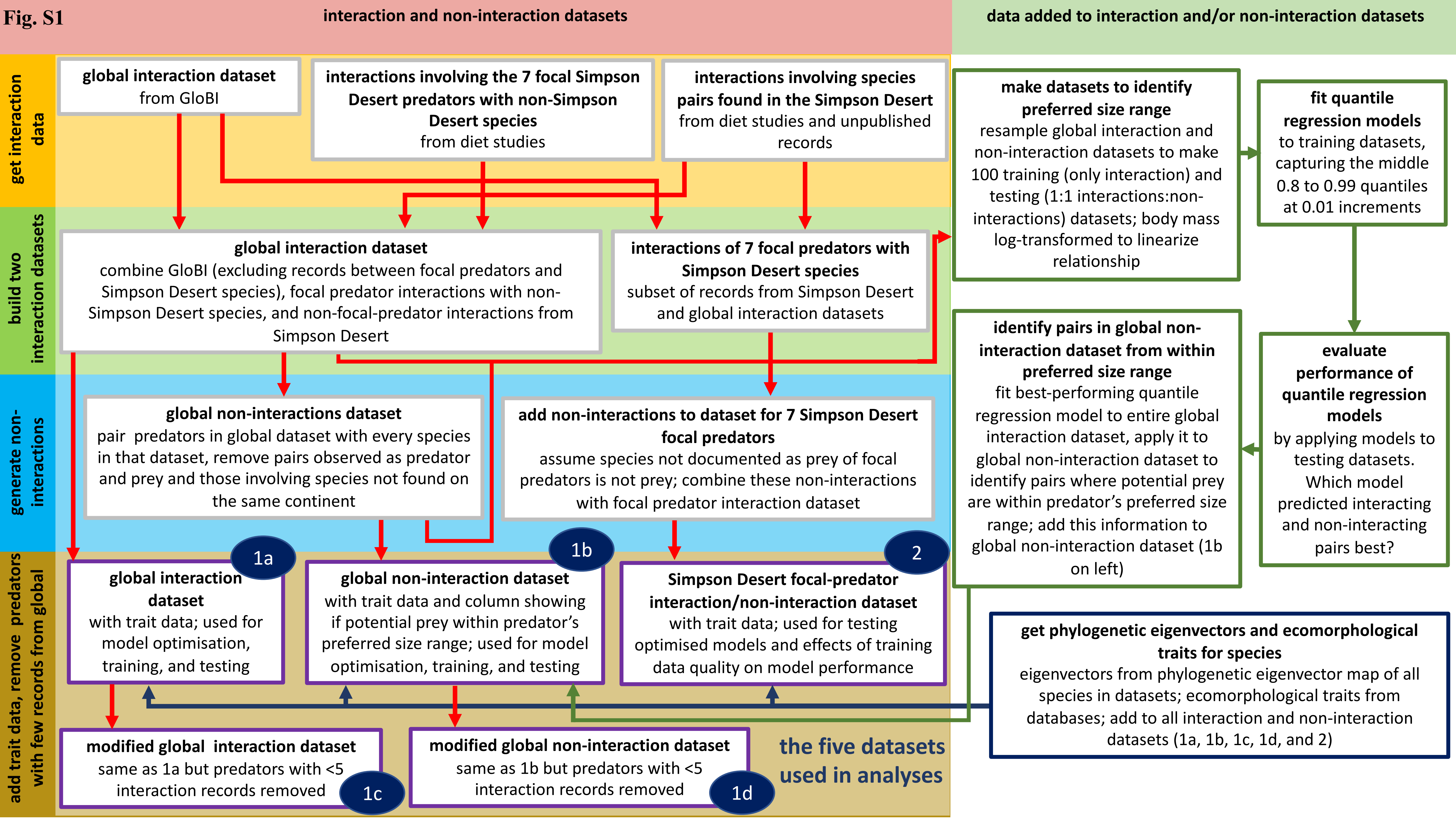
interaction and non-interaction datasets
data added to interaction and/or non-interaction datasets
get interaction data
build two interaction datasets
generate non-interactions
add trait data, remove predators with few records from global
global interaction dataset
from GloBI
interactions involving the 7 focal Simpson Desert predators with non-Simpson Desert species
from diet studies
interactions involving species pairs found in the Simpson Desert
from diet studies and unpublished records
global interaction dataset
combine GloBI (excluding records between focal predators and Simpson Desert species), focal predator interactions with non-Simpson Desert species, and non-focal-predator interactions from Simpson Desert
interactions of 7 focal predators with Simpson Desert species
subset of records from Simpson Desert and global interaction datasets
global non-interactions dataset
pair predators in global dataset with every species in that dataset, remove pairs observed as predator and prey and those involving species not found on the same continent
add non-interactions to dataset for 7 Simpson Desert focal predators
assume species not documented as prey of focal predators is not prey; combine these non-interactions with focal predator interaction dataset
1a
global interaction dataset
with trait data; used for model optimisation, training, and testing
1b
global non-interaction dataset
with trait data and column showing if potential prey within predator's preferred size range; used for model optimisation, training, and testing
2
Simpson Desert focal-predator interaction/non-interaction dataset
with trait data; used for testing optimised models and effects of training data quality on model performance
modified global interaction dataset
same as 1a but predators with <5 interaction records removed
1c
modified global non-interaction dataset
same as 1b but predators with <5 interaction records removed
1d
the five datasets used in analyses
make datasets to identify preferred size range
resample global interaction and non-interaction datasets to make 100 training (only interaction) and testing (1:1 interactions:non-interactions) datasets; body mass log-transformed to linearize relationship
fit quantile regression models
to training datasets, capturing the middle 0.8 to 0.99 quantiles at 0.01 increments
identify pairs in global non-interaction dataset from within preferred size range
fit best-performing quantile regression model to entire global interaction dataset, apply it to global non-interaction dataset to identify pairs where potential prey are within predator's preferred size range; add this information to global non-interaction dataset (1b on left)
evaluate performance of quantile regression models
by applying models to testing datasets. Which model predicted interacting and non-interacting pairs best?
get phylogenetic eigenvectors and ecomorphological traits for species
eigenvectors from phylogenetic eigenvector map of all species in datasets; ecomorphological traits from databases; add to all interaction and non-interaction datasets (1a, 1b, 1c, 1d, and 2)

Fig. S2

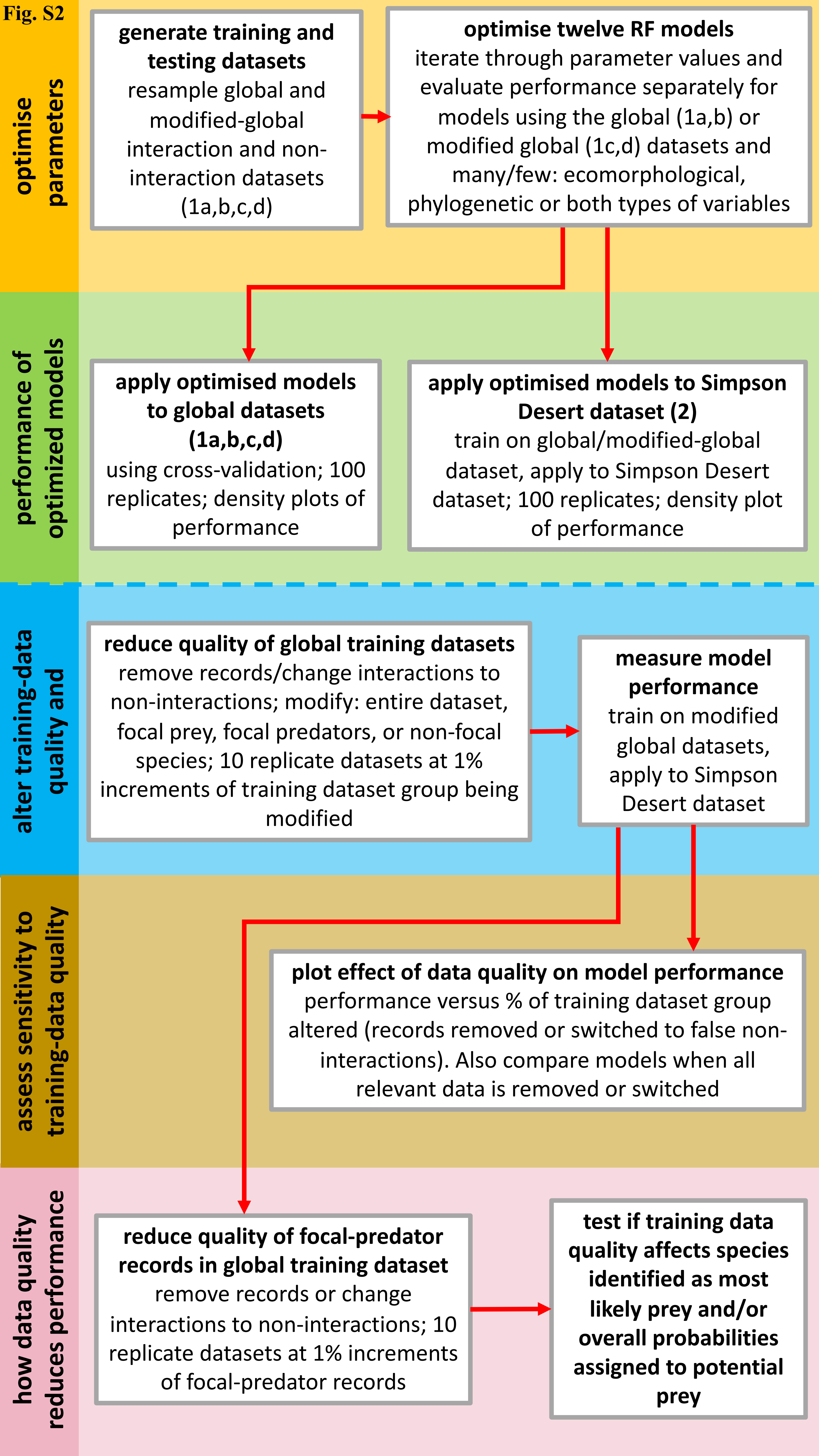
optimise parameters
generate training and testing datasets
resample global and modified-global interaction and non-interaction datasets (1a,b,c,d)
optimise twelve RF models
iterate through parameter values and evaluate performance separately for models using the global (1a,b) or modified global (1c,d) datasets and many/few: ecomorphological, phylogenetic or both types of variables
performance of optimized models
apply optimised models to global datasets (1a,b,c,d)
using cross-validation; 100 replicates; density plots of performance
apply optimised models to Simpson Desert dataset (2)
train on global/modified-global dataset, apply to Simpson Desert dataset; 100 replicates; density plot of performance
alter training-data quality and
reduce quality of global training datasets
remove records/change interactions to non-interactions; modify: entire dataset, focal prey, focal predators, or non-focal species; 10 replicate datasets at 1% increments of training dataset group being modified
measure model performance
train on modified global datasets, apply to Simpson Desert dataset
assess sensitivity to training-data quality
plot effect of data quality on model performance
performance versus % of training dataset group altered (records removed or switched to false non-interactions). Also compare models when all relevant data is removed or switched
how data quality reduces performance
reduce quality of focal-predator records in global training dataset
remove records or change interactions to non-interactions; 10 replicate datasets at 1% increments of focal-predator records
test if training data quality affects species identified as most likely prey and/or overall probabilities assigned to potential prey

**Supplementary Table S1 – Ecomorphological variables**

| **ecomorphological variable** | **data type** | **for which animals** |
| --- | --- | --- |
| body mass* | continuous | birds and mammals |
| nocturnal | binary | birds and mammals |
| diurnal | binary | birds and mammals |
| crepuscular | binary | birds and mammals |
| ability to fly | binary | birds and mammals |
| consumes plants | binary | birds and mammals |
| consumes vertebrates | binary | birds and mammals |
| consumes invertebrates | binary | birds and mammals |
| strata use – below water | percent | birds |
| strata use – water surface | percent | birds |
| strata use - ground | percent | birds |
| strata use - understory | percent | birds |
| strata use – mid-high | percent | birds |
| strata use - canopy | percent | birds |
| strata use - aerial | percent | birds |
| strata use - pelagic | percent | birds |
| strata use - marine | percent | mammals |
| strata use - ground | percent | mammals |
| strata use - scansorial | percent | mammals |
| strata use – arboreal | percent | mammals |
| strata use - aerial | percent | mammals |

*body mass ranged from 2.13 to 30,000000 g

**Supplementary Table S2 – Datasets/models and what they were used for**

| data/model | | | uses/analyses | | |
|---|---|---|---|---|---|
| **training data** | **testing data** | **variables** | **optimise parameters** | **performance (Fig. 1)** | **effect of data quality on performance (Fig. 3 & 4)** |
| global | global | many, both types | ✓ | ✓ | ✗ |
| global | global | few, both types | ✓ | ✓ | ✗ |
| global | global | many, phylogenetic eigenv. | ✓ | ✓ | ✗ |
| global | global | few, phylogenetic eigenv. | ✓ | ✓ | ✗ |
| global | global | many, ecomorphological | ✓ | ✓ | ✗ |
| global | global | few, ecomorphological | ✓ | ✓ | ✗ |
| modified global | modified global | many, both types | ✓ | ✓ | ✗ |
| modified global | modified global | few, both types | ✓ | ✓ | ✗ |
| modified global | modified global | many, phylogenetic eigenv. | ✓ | ✓ | ✗ |
| modified global | modified global | few, phylogenetic eigenv. | ✓ | ✓ | ✗ |
| modified global | modified global | many, ecomorphological | ✓ | ✓ | ✗ |
| modified global | modified global | few, ecomorphological | ✓ | ✓ | ✗ |
| global | Simpson Desert | many, both types | ✓ | ✓ | ✗ |
| global | Simpson Desert | few, both types | ✗ | ✓ | ✗ |
| global | Simpson Desert | many, phylogenetic eigenv. | ✗ | ✓ | ✗ |
| global | Simpson Desert | few, phylogenetic eigenv. | ✗ | ✓ | ✗ |
| global | Simpson Desert | many, ecomorphological | ✗ | ✓ | ✗ |
| global | Simpson Desert | few, ecomorphological | ✗ | ✓ | ✗ |
| modified global | Simpson Desert | many, both types | ✗ | ✓ | ✓ |
| modified global | Simpson Desert | few, both types | ✗ | ✓ | ✗ |
| modified global | Simpson Desert | many, phylogenetic eigenv. | ✗ | ✓ | ✗ |
| modified global | Simpson Desert | few, phylogenetic eigenv. | ✗ | ✓ | ✗ |
| modified global | Simpson Desert | many, ecomorphological | ✗ | ✓ | ✗ |
| modified global | Simpson Desert | few, ecomorphological | ✗ | ✓ | ✗ |

**Supplementary Table S3 – Different models and their performance according to different metrics**

The first twelve rows show performance of the models on global datasets (original and modified), while the next twelve rows show performance on the Simpson Desert dataset when models were training on either the original or modified global dataset. The best performing model according to each measure in each set is highlighted.

| model input and test data | | | | performance metrics | | | | |
|---|---|---|---|---|---|---|---|---|
| **dataset** | **full/modified global dataset** | **many/few variables** | **variables** | **true skill statistic** | **accuracy** | **Matthew's correlation coefficient** | **sensitivity/true positive rate** | **specificity** |
| global | full | many | both types | 0.767 ± 0.015 | 0.884 ± 0.007 | 0.767 ± 0.015 | 0.879 ± 0.011 | 0.889 ± 0.011 |
| global | full | few | both types | 0.754 ± 0.015 | 0.877 ± 0.007 | 0.756 ± 0.015 | 0.839 ± 0.011 | 0.915 ± 0.009 |
| global | modified | many | both types | 0.636 ± 0.019 | 0.818 ± 0.009 | 0.637 ± 0.018 | 0.845 ± 0.012 | 0.791 ± 0.015 |
| global | modified | few | both types | 0.619 ± 0.020 | 0.810 ± 0.010 | 0.621 ± 0.020 | 0.844 ± 0.014 | 0.775 ± 0.016 |
| global | full | many | ecomorphological | 0.754 ± 0.016 | 0.877 ± 0.008 | 0.755 ± 0.016 | 0.851 ± 0.013 | 0.903 ± 0.010 |
| global | full | few | ecomorphological | 0.670 ± 0.017 | 0.835 ± 0.009 | 0.671 ± 0.017 | 0.833± 0.012 | 0.837 ± 0.014 |
| global | modified | many | ecomorphological | 0.597 ± 0.015 | 0.799 ± 0.008 | 0.604 ± 0.015 | 0.871 ± 0.011 | 0.727 ± 0.014 |
| global | modified | few | ecomorphological | 0.533 ± 0.020 | 0.766 ± 0.010 | 0.534 ± 0.020 | 0.800 ± 0.014 | 0.733 ± 0.016 |
| global | full | many | phylogenetic eigenvectors | 0.741 ± 0.018 | 0.870 ± 0.009 | 0.741 ± 0.018 | 0.860 ± 0.013 | 0.880 ± 0.013 |
| global | full | few | phylogenetic eigenvectors | 0.724 ± 0.016 | 0.862 ± 0.008 | 0.724 ± 0.016 | 0.861 ± 0.012 | 0.863 ± 0.012 |
| global | modified | many | phylogenetic eigenvectors | 0.596 ± 0.018 | 0.798 ± 0.009 | 0.597 ± 0.018 | 0.810 ± 0.014 | 0.787 ± 0.014 |
| global | modified | few | phylogenetic eigenvectors | 0.593 ± 0.018 | 0.796 ± 0.009 | 0.594 ± 0.018 | 0.830 ± 0.013 | 0.763 ± 0.014 |
| Simpson Desert | full | many | both types | 0.279 ± 0.023 | 0.501 ± 0.02 | 0.259 ± 0.018 | 0.906 ± 0.015 | 0.373 ± 0.029 |
| Simpson Desert | full | few | both types | 0.346 ± 0.02 | 0.593 ± 0.015 | 0.298 ± 0.016 | 0.826 ± 0.014 | 0.519 ± 0.021 |
| Simpson Desert | modified | many | both types | 0.452 ± 0.018 | 0.669 ± 0.011 | 0.386 ± 0.016 | 0.835 ± 0.02 | 0.617 ± 0.017 |
| Simpson Desert | modified | few | both types | 0.434 ± 0.023 | 0.647 ± 0.016 | 0.371 ± 0.019 | 0.851 ± 0.02 | 0.583 ± 0.022 |
| Simpson Desert | full | many | ecomorphological | 0.382 ± 0.014 | 0.606 ± 0.008 | 0.328 ± 0.012 | 0.853 ± 0.014 | 0.529 ± 0.012 |
| Simpson Desert | full | few | ecomorphological | 0.25 ± 0.013 | 0.478 ± 0.012 | 0.237 ± 0.012 | 0.907 ± 0.017 | 0.343 ± 0.02 |
| Simpson Desert | modified | many | ecomorphological | 0.402 ± 0.016 | 0.637 ± 0.012 | 0.344 ± 0.013 | 0.825 ± 0.014 | 0.578 ± 0.018 |
| Simpson Desert | modified | few | ecomorphological | 0.251 ± 0.016 | 0.502 ± 0.012 | 0.228 ± 0.013 | 0.863 ± 0.016 | 0.388 ± 0.018 |
| Simpson Desert | full | many | phylogenetic eigenvectors | 0.282 ± 0.026 | 0.516 ± 0.022 | 0.255 ± 0.02 | 0.88 ± 0.018 | 0.402 ± 0.031 |
| Simpson Desert | full | few | phylogenetic eigenvectors | 0.28 ± 0.029 | 0.509 ± 0.023 | 0.256 ± 0.023 | 0.892 ± 0.014 | 0.388 ± 0.031 |
| Simpson Desert | modified | many | phylogenetic eigenvectors | 0.396 ± 0.022 | 0.67 ± 0.011 | 0.34 ± 0.019 | 0.751 ± 0.022 | 0.644 ± 0.016 |
| Simpson Desert | modified | few | phylogenetic eigenvectors | 0.398 ± 0.024 | 0.633 ± 0.018 | 0.34 ± 0.02 | 0.826 ± 0.014 | 0.572 ± 0.024 |

### Supplementary Table S4 – Retained variables in the few-variable models

| **full/modified global dataset** | **variables** | **variables retained** | | | | | | | | | |
|---|---|---|---|---|---|---|---|---|---|---|---|
| modified | both types | source bodymass | source flight | target bodymass | target flight | source forstrat understory | source forstrat midhigh | source forstrat aerial | target forstrat understory | target forstrat midhigh | target forstrat aerial |
| full | both types | source bodymass | source flight | source eat_invs | target bodymass | target flight | target eat_invs | source forstrat understory | source forstrat midhigh | target forstrat understory | target forstrat midhigh |
| modified | ecomorphological | source bodymass | source eat_invs | target bodymass | target eat_invs | source forstrat understory | source forstrat midhigh | source forstrat canopy | target forstrat understory | target forstrat midhigh | target forstrat canopy |
| full | ecomorphological | source bodymass | source eat_plants | target bodymass | target eat_plants | source forstrat understory | source forstrat midhigh | source forstrat canopy | target forstrat understory | target forstrat midhigh | target forstrat canopy |
| modified | phylogenetic eigenvectors | target eig1 | target eig2 | target eig3 | target eig4 | target eig5 | source eig1 | source eig2 | source eig3 | source eig4 | source eig5 |
| full | phylogenetic eigenvectors | target eig2 | target eig3 | target eig4 | target eig5 | target eig6 | source eig2 | source eig3 | source eig4 | source eig5 | source eig6 |

| **full/modified global dataset** | **variables** | **variables retained** | | | | | | | | | |
|---|---|---|---|---|---|---|---|---|---|---|---|
| modified | both types | target eig1 | target eig2 | target eig3 | target eig4 | target eig5 | source eig1 | source eig2 | source eig3 | source eig4 | source eig5 |
| full | both types | target eig1 | target eig2 | target eig3 | target eig4 | target eig5 | source eig1 | source eig2 | source eig3 | source eig4 | source eig5 |
| modified | ecomorphological | NA | NA | NA | NA | NA | NA | NA | NA | NA | NA |
| full | ecomorphological | NA | NA | NA | NA | NA | NA | NA | NA | NA | NA |
| modified | phylogenetic eigenvectors | NA | NA | NA | NA | NA | NA | NA | NA | NA | NA |
| full | phylogenetic eigenvectors | NA | NA | NA | NA | NA | NA | NA | NA | NA | NA |

*eig = phylogenetic eigenvector; ForStrat = forest strata/habitat use; eat_invs = diet includes invertebrates; eat_plant = diet includes plants; source = predator; target = prey

**Supplementary Table S5 - performance of models trained on weighted *versus* unweighted records**

Interaction and non-interaction records in train data were either (1) weighted so their sum-total weights were equal, or (2) were not weighted.

| | **unweighted** | | **weighted** | |
|---|---|---|---|---|
| | **mean** | **sd** | **mean** | **sd** |
| **Global dataset** | 0.690 | 0.018 | 0.701 | 0.017 |
| **Simpson Desert dataset** | 0.355 | 0.028 | 0.376 | 0.024 |

**Table S6 — Optimised parameters for the different models**

| model input | | | model parameters | | | | | |
|---|---|---|---|---|---|---|---|---|
| **full/modified global dataset** | **many/few variables** | **variables** | **non-interactions**[1] | **inside**[2] | **mtry**[3] | **threshold**[4] | **ntrees**[5] | **depth**[6] |
| full | many | both | 3 | 0.5 | 4 | 0.41 | 600 | 30 |
| modified | many | both | 4.75 | 2.5 | 42 | 0.31 | 400 | 0 |
| full | few | both | 4.5 | 0.5 | 5 | 0.49 | 200 | 20 |
| modified | few | both | 3.5 | 1.75 | 8 | 0.36 | 350 | 1000 |
| full | many | ecomorphological | 4 | 0.75 | 11 | 0.42 | 650 | 500 |
| modified | many | ecomorphological | 3 | 1.5 | 23 | 0.34 | 250 | 100 |
| full | few | ecomorphological | 4.75 | 3.25 | 6 | 0.35 | 650 | 500 |
| modified | few | ecomorphological | 4.75 | 3.5 | 4 | 0.39 | 250 | 100 |
| full | many | phylogenetic eigenv. | 1.5 | 2 | 3 | 0.44 | 750 | 20 |
| modified | many | phylogenetic eigenv. | 4.75 | 1 | 8 | 0.36 | 350 | 1000 |
| full | few | phylogenetic eigenv. | 3 | 0.5 | 3 | 0.41 | 600 | 30 |
| modified | few | phylogenetic eigenv. | 4 | 2.25 | 2 | 0.36 | 350 | 1000 |

[1]**non-interactions** = number of non-interactions relative to number of interactions in the training dataset. [2]**inside** = number of non-interactions from inside the preferred size range relative to number of non-interactions from outside the preferred size range. [3]**mtry** = number of traits/variables (randomly sampled from possible variables) considered at each split in the decision trees. [4]**threshold** = probability threshold for assigning species pairs as interacting (predator and prey) or not. [5]**ntrees** = number of decision trees in the random forest. [6]**depth** = maximum depth of decision trees in the random forest.
